# State-dependent pontine ensemble dynamics and interactions with cortex across sleep states

**DOI:** 10.1101/752683

**Authors:** Tomomi Tsunematsu, Amisha A Patel, Arno Onken, Shuzo Sakata

## Abstract

The pontine nuclei play a crucial role in sleep-wake regulation. However, pontine ensemble dynamics underlying sleep regulation remain poorly understood. By monitoring population activity in multiple pontine and adjacent brainstem areas, here we show slow, state-predictive pontine ensemble dynamics and state-dependent interactions between the pons and the cortex in mice. On a timescale of seconds to minutes, pontine populations exhibit diverse firing across vigilance states, with some of these dynamics being attributed to cell type-specific activity. Pontine population activity can predict pupil dilation and vigilance states: pontine neurons exhibit longer predictable power compared with hippocampal neurons. On a timescale of sub-seconds, pontine waves (P-waves) are observed as synchronous firing of pontine neurons primarily during rapid eye movement (REM) sleep, but also during non-REM (NREM) sleep. Crucially, P-waves functionally interact with cortical activity in a state-dependent manner: during NREM sleep, hippocampal sharp wave-ripples (SWRs) precede P-waves. On the other hand, P-waves during REM sleep are phase-locked with ongoing hippocampal theta oscillations and are followed by burst firing in a subset of hippocampal neurons. Thus, the directionality of functional interactions between the hippocampus and pons changes depending on sleep states. This state-dependent global coordination between pontine and cortical regions implicates distinct functional roles of sleep.

## Introduction

The sleep-wake cycle is a fundamental homeostatic process across animal species (Anafi et al., 2019; Aulsebrook et al., 2016; Siegel, 2005). In addition to the physiological functions of sleep (Boyce et al., 2017; Brown et al., 2012; Imeri and Opp, 2009; Liu and Dan, 2019; Rasch and Born, 2013; Sara, 2017; Siegel, 2005; Stickgold et al., 2001; Tononi and Cirelli, 2014), the abnormalities in the sleep-wake cycle are associated with various diseases and disorders (Brown et al., 2012; Irwin, 2015; Mander et al., 2017; Musiek and Holtzman, 2016).

Sleep states are typically classified into two major states, non-rapid eye movement (NREM) sleep and REM sleep. While numerous brain regions and cell-types have been identified as part of sleep-regulating circuits (Adamantidis et al., 2007; Brown et al., 2012; Herice et al., 2019; Jouvet, 1962; Luppi et al., 2017; Moruzzi, 1963; Peever and Fuller, 2017; Scammell et al., 2017; Tsunematsu et al., 2014; Weber et al., 2015; Weber and Dan, 2016; Zhang et al., 2019), sleep-related neural firing and oscillations have also been described across cortical and subcortical regions (Brown et al., 2012; Buzsaki, 2015; Herice et al., 2019; Hobson et al., 1975; Liu and Dan, 2019; McCarley and Hobson, 1971; Rasch and Born, 2013; Sakai, 1985; Scammell et al., 2017; Steriade, 2006; Weber et al., 2015; Weber et al., 2018). For example, cortical slow oscillations, sleep spindles and hippocampal sharp wave-ripples (SWRs) are prominent neural events during NREM sleep whereas theta oscillations and ponto-geniculo-occipital (PGO) or pontine (P) waves are seen during REM sleep (Bizzi and Brooks, 1963; Buzsaki, 2002, 2015; Callaway et al., 1987; Datta, 1997; Jouvet, 1969; Montgomery et al., 2008; Rasch and Born, 2013; Steriade, 2006; Steriade et al., 1993b). Although neural ensemble dynamics underlying these sleep-related neural events in the cortex and the thalamus have been well described (Buzsaki, 2002, 2015; Steriade, 2006; Steriade et al., 1993a), little is known about population activity within the brainstem. Toward a better understanding of functional roles of sleep states, it is essential to characterize state-dependent changes in brainstem network activity and their functional interactions with cortical regions across sleep states.

The brainstem, including the midbrain, pons and medulla has long been implicated in the sleep-wake cycle (Brown et al., 2012; Herice et al., 2019; Jouvet, 1962; Liu and Dan, 2019; Luppi et al., 2017; Rasch and Born, 2013; Saper et al., 2010; Scammell et al., 2017; Weber et al., 2015; Weber and Dan, 2016). It contains various nuclei, each of which consists of diverse cell-types and exhibits state-dependent firing (Brown et al., 2012; Herice et al., 2019; Liu and Dan, 2019; Luppi et al., 2017; Rasch and Born, 2013; Scammell et al., 2017; Weber et al., 2015; Weber and Dan, 2016; Weber et al., 2018; Zhang et al., 2019). However, it remains poorly explored how brainstem populations act in concert. For example, it is still unclear to what extent their activity exhibits anticipatory dynamics for ongoing vigilant states. In addition, it is also unclear whether and how brainstem populations functionally interact with various neural oscillations or events in the cortex across sleep states. Characterizing these physiological properties is crucial to uncover the roles of brainstem populations in sleep regulation and ultimately the functions of sleep states.

In the present study, we adopt several *in vivo* electrophysiological approaches in mice to investigate state-dependent ensemble dynamics in the brainstem, mainly the pons. We show that on a timescale of seconds to minutes, pontine neurons show state-dependent firing with cell type-specificity. They also have a longer predictive power for vigilance states compared to those in the hippocampus. On a timescale of sub-seconds, we find state-dependent functional interactions between the pons and the cortex, with a focus on P-waves: during NREM sleep, the timing of P-waves is phase-locked with various cortical oscillations and hippocampal SWRs precede P-waves. During REM sleep, P-waves co-occur with hippocampal theta and precede burst firing of hippocampal neurons. These results imply that pontine populations not only play a regulatory role in the sleep-wake cycle, but also contribute to global state-dependent dynamics across brain regions.

## Results

### Brainstem population recording across sleep-wake cycles

To investigate the state-dependency of brainstem population activity, we inserted a silicon probe into the mouse brainstem in a head-fixed condition, together with simultaneous monitoring of cortical electroencephalograms (EEGs), electromyograms (EMGs) and pupil dilation (**Fig. 1**). Recorded regions spanned across multiple nuclei, including the sublaterodorsal nucleus, pontine reticular nucleus, medial preoptic nucleus, parabrachial nucleus, pontine central gray, laterodorsal tegmental nucleus and other surrounding areas according to post-mortem histological analysis (**Supplementary Fig. 1**). Although a majority of neurons were recorded from the pons, we refer to recorded populations as “brainstem” neurons because some cells were located in the midbrain and medulla, but not the hypothalamus.

**Figure 1.**
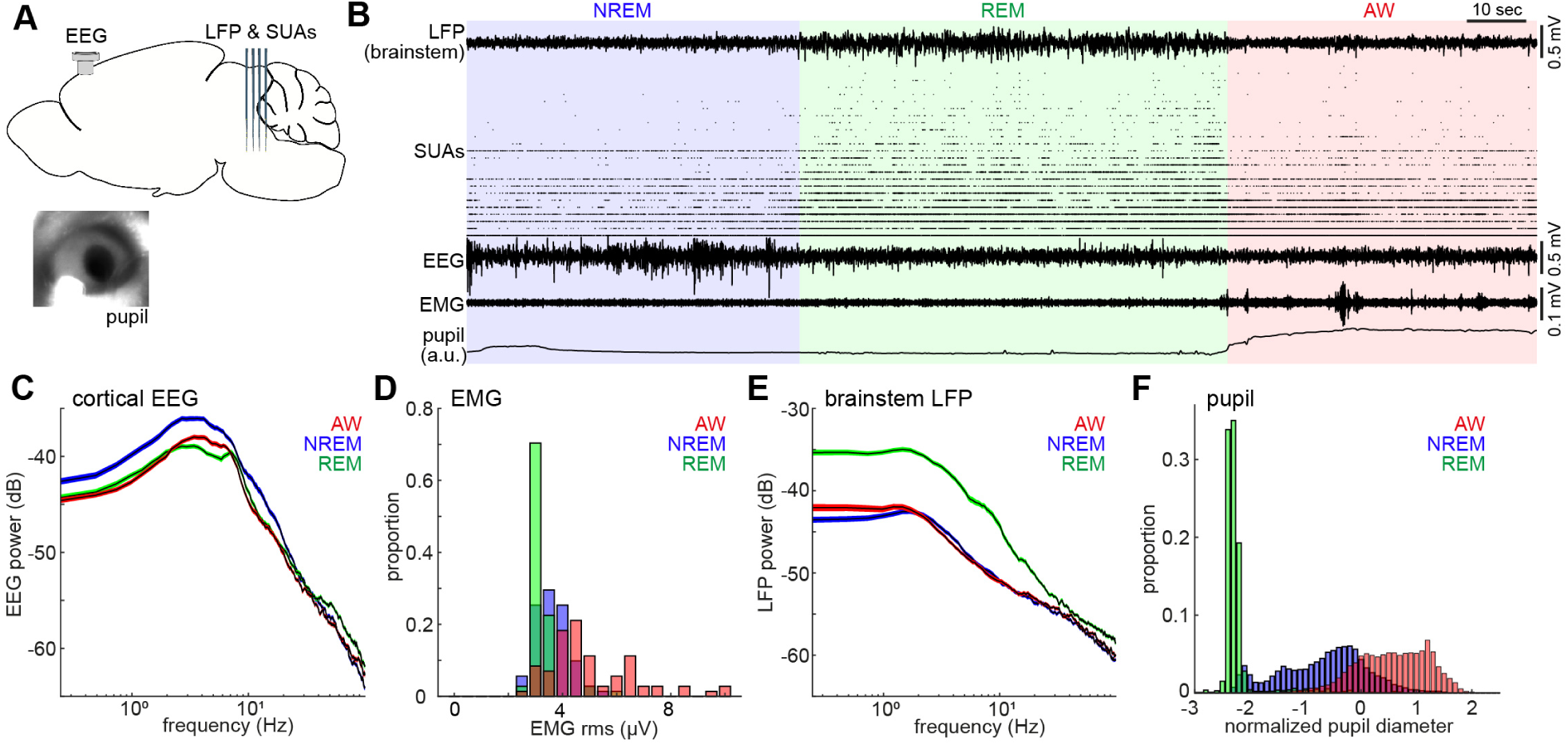
Population activity in the brainstem across the sleep-wake cycle. **A**. A diagram of experimental approaches, showing the insertion of a silicon probe for extracellular recording in the brainstem and a screw for cortical EEG recording. Pupil dilation and EMGs were also monitored in a head-fixed condition. **B**. An example of multiple electrophysiological readings across three behavioral states, including local field potentials (LFPs) in the brainstem (locally subtracted LFP signals), brainstem single unit activities (SUAs), cortical EEG, EMG and normalized pupil diameter. REM, rapid eye movement sleep; NREM, non-REM sleep; AW, wakefulness. **C and E**. Power spectrum density of cortical EEGs (**C**) and brainstem LFPs (**E**) across three behavioral states. Spectrum was computed during every 4-sec window. Errors indicate SEM. **D and F**. Distribution of EMG signals (root mean square) (**D**) and normalized pupil diameter (**F**) across three behavioral states. Pupil diameter was normalized as z-score.

The sleep-wake cycle was classified based on cortical EEGs and EMGs in every 4 second. Based on the classified states, we observed clear state-dependency across measurements (**Figs. 1C-F**): wakefulness was characterized by high muscle tone (**Fig. 1D**) and pupil dilation (**Fig. 1F**) whereas NREM sleep was characterized by higher power of slow oscillations (**Fig. 1C**) and a wider dynamic range of pupil diameter (**Fig. 1F**). REM sleep was distinct from the other states, with respect to prominent theta oscillations (**Fig. 1C**), low muscle tone (**Fig. 1D**), higher brainstem LFPs power (**Fig. 1E**) and fully constricted pupil (**Fig. 1F**). The higher power of brainstem LFPs during REM sleep was preserved across animals (7 animals, 9 recordings) (**Supplementary Fig. 2**).

Neuronal spiking activity in the brainstem also demonstrated rich state-dependent properties (**Fig. 1B**). For example, a subset of neurons fired exclusively during REM sleep, indicating state dependent population firing on a timescale of second-to-minute. In addition, we also observed frequent burst firing across neurons on a sub-second timescale during REM sleep. In the following analysis, we investigate state-dependent brainstem neural ensembles on two distinct timescales: a long timescale of seconds to minutes (**Figs. 2-4**) and a short sub-second timescale (**Figs. 5-7**).

**Figure 2.**
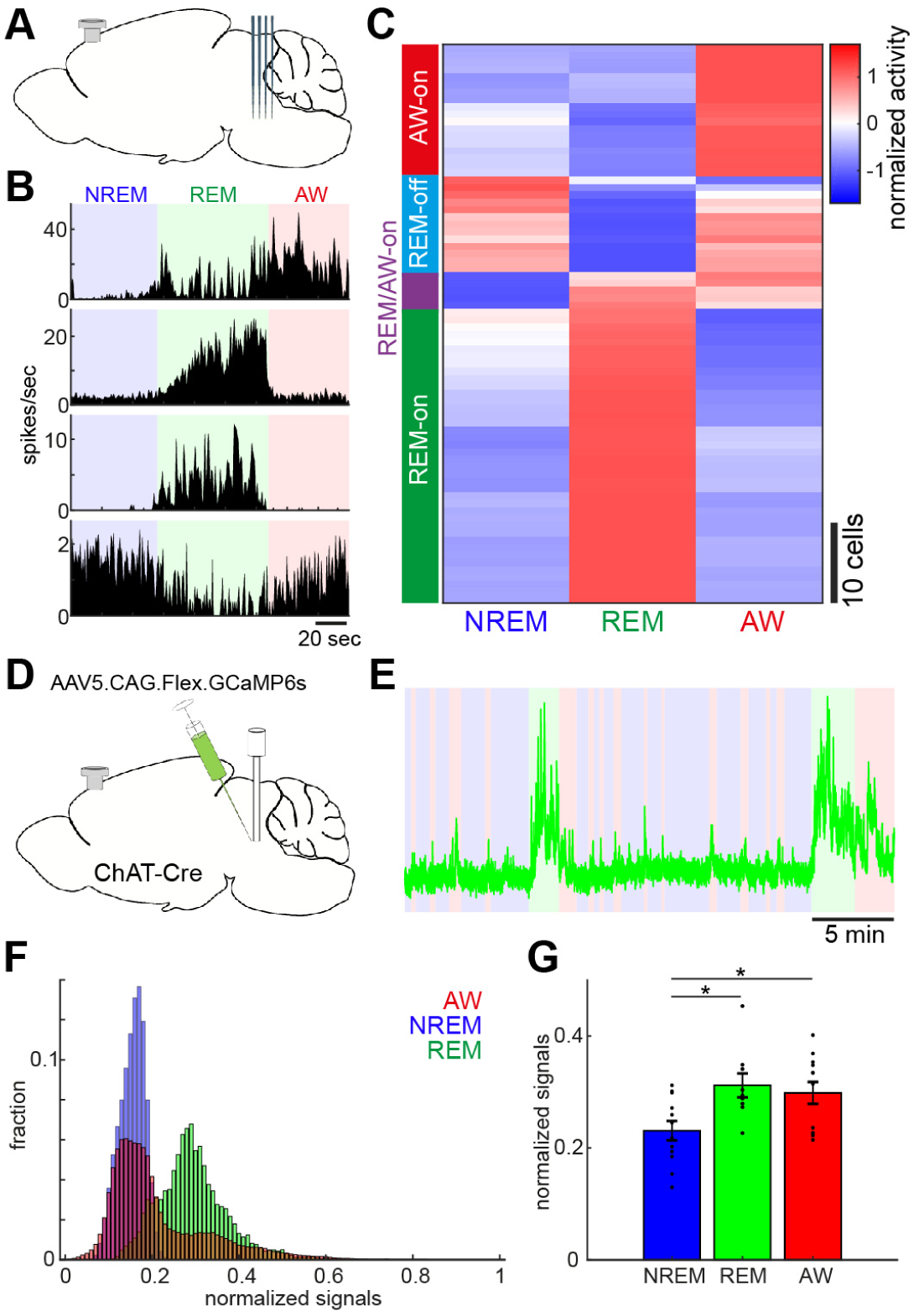
Diverse and cell-type-specific state-dependent firing in brainstem neurons. **A**. A diagram of an experimental approach, showing a silicon probe and a cortical EEG electrode. **B**. Four examples of simultaneously recording neurons. **C**. Classification of functional classes. Firing rates across three behavioral states were normalized as z-score for individual cells, then a hierarchical clustering was applied. **D**. A diagram of an experimental approach for fiber photometry-based Ca^2+^ imaging from pontine cholinergic neural populations in a freely behaving condition. **E**. An example of fluorescent signals across sleep-wake cycles. Fluorescent signals (470 nm) were normalized by off-peak (405 nm) signals. red, wakefulness, blue, NREM sleep, green, REM sleep. **F**. Distributions of fluorescent signals across three behavioral states. **G**. Group statistics of average signals from 12 recordings from 4 mice (F_2,32_ = 5.12, *p* = 0.012, one-way ANOVA). *, *p* < 0.05 with post-hoc Tukey’s honest significant difference criterion.

**Figure 3.**
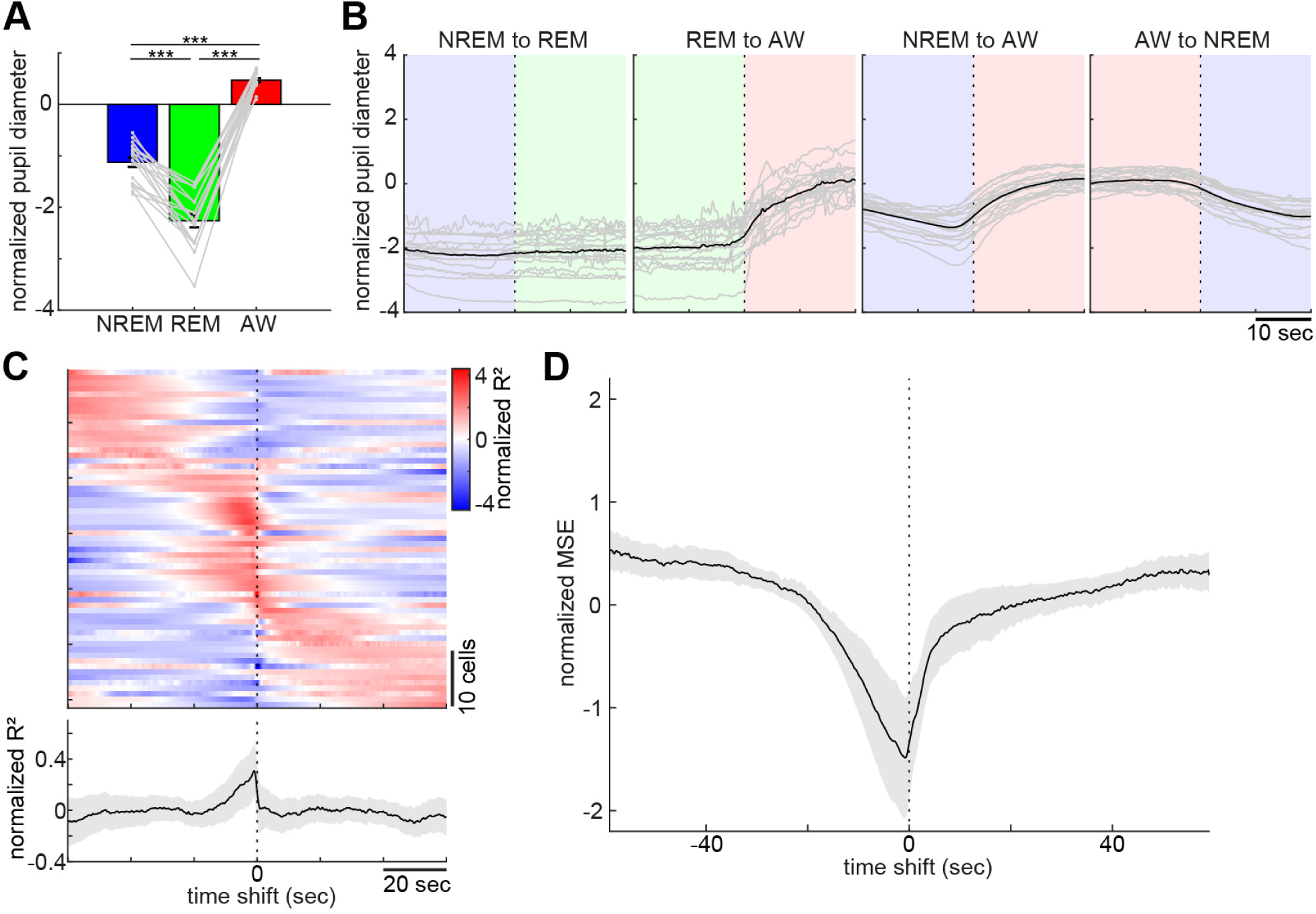
Pupil dilation across the sleep-wake cycle and prediction of pupil dilation by brainstem populations. **A**. Mean normalized (z-scored) pupil diameter across the sleep-wake cycle (n = 18, F_2, 53_ = 220.33, *p* < 0.0001. one-way ANOVA). ***, *p* < 0.0001, with post-hoc Tukey’s honest significant difference criterion. **B**. Pupil dilation at the transition of behavioral states (n = 18). **C**. Linear regression analysis to predict pupil diameter by individual neuronal activity. (*top*) Normalized (z-scored) cross-validated R^2^ values are color-coded across neurons. (*bottom*) The average of normalized cross-validated R^2^ values. Error, SEM. **D**. Multiple linear regression analysis to predict pupil diameter by simultaneously recorded neuronal activity. The average of normalized (z-scored) root mean square errors across datasets (n = 6). Error, SEM.

**Figure 4.**
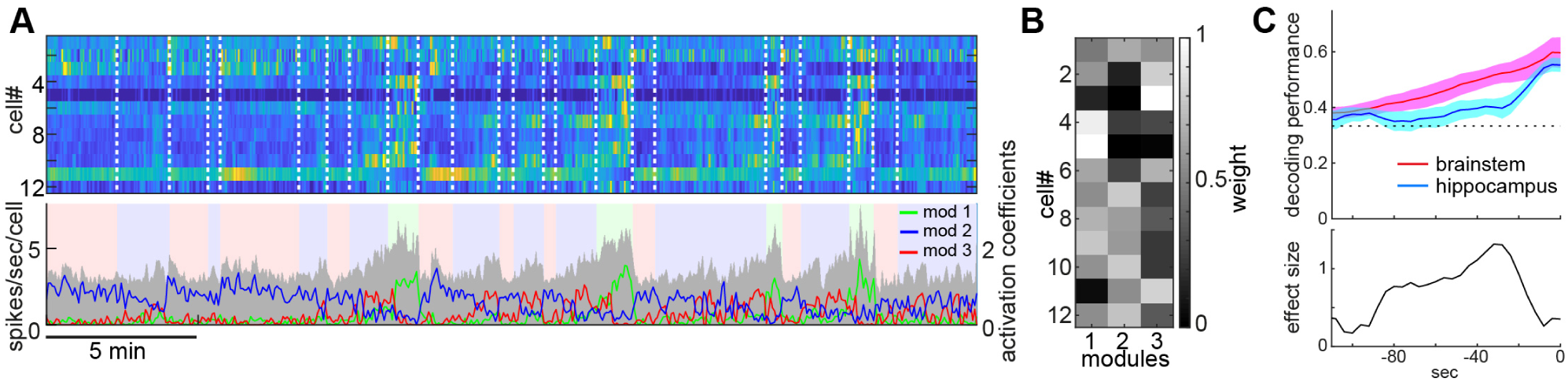
State-dependent brainstem population dynamics and their predictability for behavioral states. **A**. Simultaneously recorded brainstem neurons across states and modules extracted by non-negative matrix factorization (NMF). (top) Firing profiles of brainstem neurons. Firing rate was normalized by the maximum firing rate and the normalized values were color-coded. Dotted lines indicate the timing of state transitions. (bottom) Average population activity (gray) and activation coefficients for each module derived by NMF. Background colors indicate behavioral states (red, AW; blue, NREM; green, REM).. **B**. Weights across neurons for each module. **C**. Decoding of behavioral states from population activity. *top*, Decoding performance of brainstem and hippocampal neural populations for behavioral states as a function of time-shift. Error, SEM. *bottom*, Effect size as a function of time-shift.

**Figure 5.**
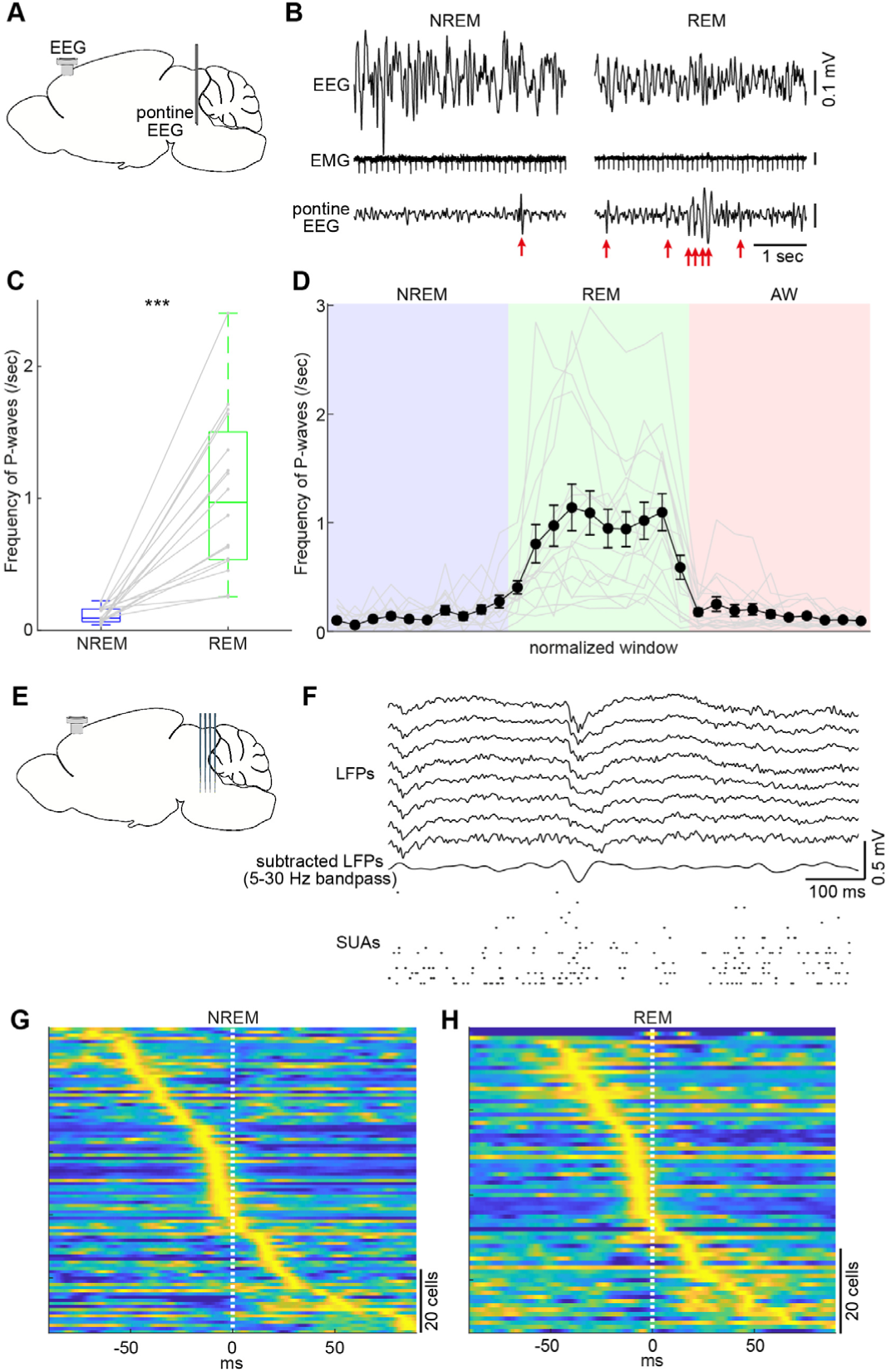
Pontine waves (P-waves) in the mouse. **A**. A diagram of an experimental approach, with showing a bipolar electrode in the pons and a cortical EEG electrode. **B**. Examples of P-waves during NREM (left) and REM sleep (right), with showing cortical EEG and EMG traces. **C**. Frequency of P-waves during NREM and REM sleep (n = 16). ***, *p* < 0.0001, two-tailed *t*-test. **D**. Temporal evolution of P-wave frequency. Duration of each state episode was normalized to one. Error, SEM. **E**. A diagram of the experimental approach showing a multi-shank silicon probe and cortical EEG electrode. **F**. An example of P-wave, showing LFPs from a shank, filtered, subtracted LFPs, and multiple single unit activity. **G and H**. Pooled peri-event time histograms of brainstem single units relative to P-wave timing during NREM (**G**) and REM sleep (**H**). Time zero is the timing of P-waves (trough time). Each peri-event time histogram is color-coded by normalizing the maximum firing rate for each cell. The order of single units was sorted by the peak timing in each state.

**Figure 6.**
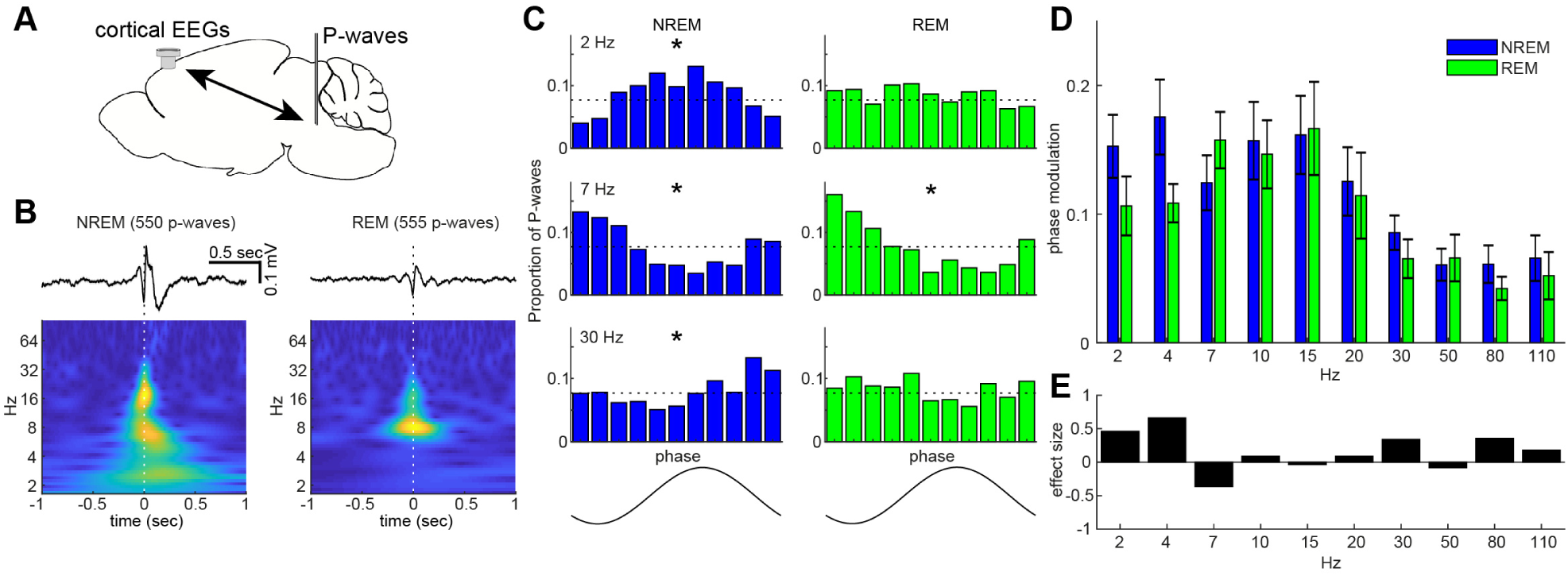
State-dependent interactions between P-waves and cortical oscillations. **A**. A diagram of an experimental approachshowing a bipolar electrode in the pons and cortical EEG electrode. **B**. Examples of averaged event-triggered cortical EEGs and scalograms during NREM (left) and REM sleep (right). Time zero is the timing of P-waves (trough time). **C**. Examples of phase-histograms. Cortical EEGs were filtered at certain frequency bands and the proportion of P-waves elicited in each phase bin was calculated. *, *p* < 0. 01, Rayleigh’s test. **D**. Phase modulations across frequency bands of cortical EEGs. The phase modulation index was defined as the proportion in the preferred bin (the bin with maximal percentage) minus that in the opposite bin (the bin 180° apart). Error, SEM. **E**. Effect size of states across frequency bands.

**Figure 7.**
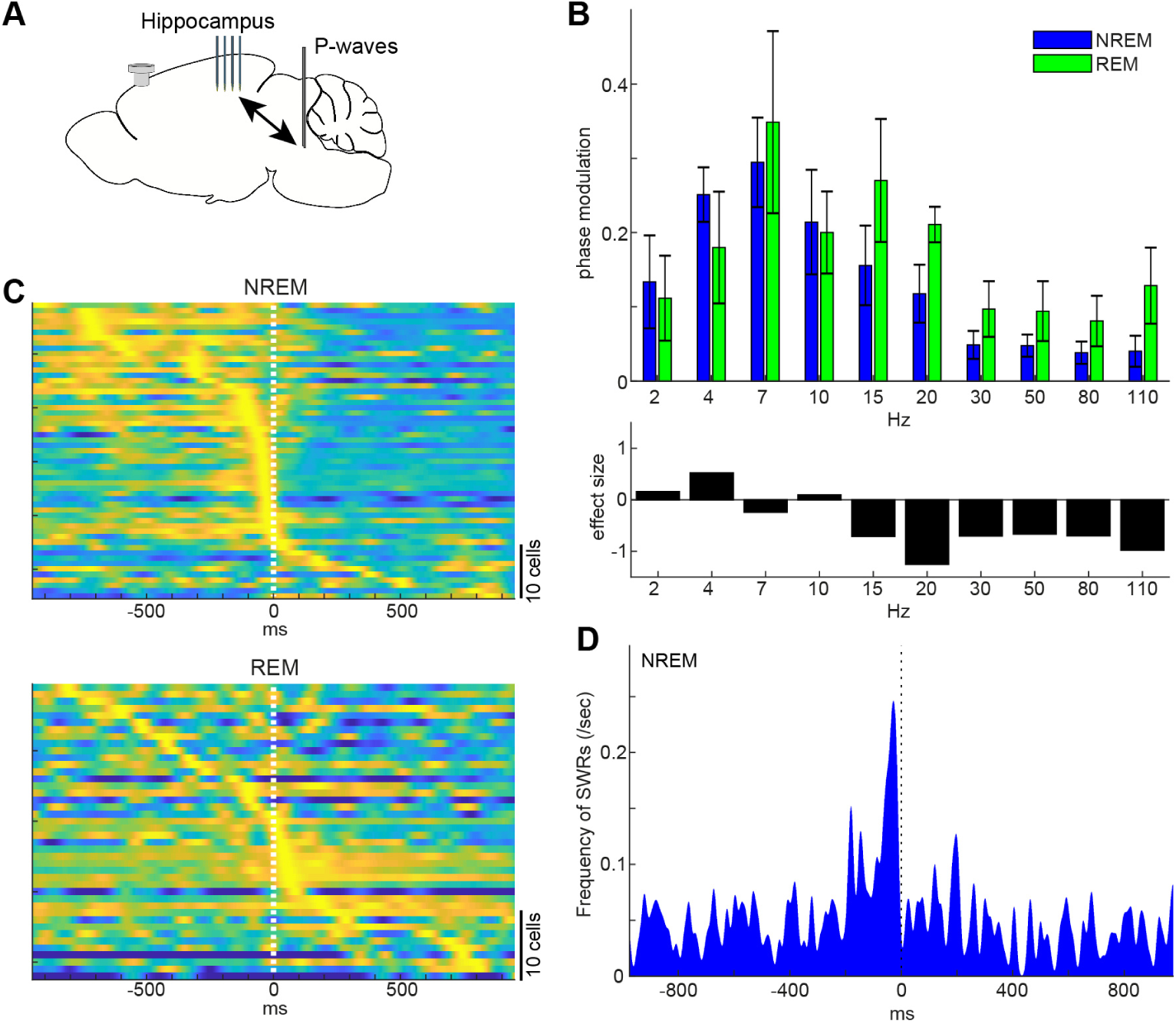
State-dependent interactions between P-waves and hippocampal activity. **A**. A diagram of an experimental approach showing a multi-shank silicon probe in the hippocampus, a bipolar electrode in the pons and a cortical EEG electrode. **B**. Phase modulations across frequency bands of hippocampal LFPs (top) and effect size of states (bottom). Error, SEM. **C**. Pooled peri-event time histograms of hippocampal single units relative to P-wave timing in NREM (top) and REM sleep (bottom). Time zero is the timing of P-waves (trough time). Each peri-event time histogram is color-coded by normalizing the maximum firing rate for each cell. The order of single units was sorted by the peak timing in each state. **D**. Frequency of hippocampal SWRs relative to P-wave timing in NREM sleep. Time zero is the timing of P-waves (trough time).

### Diversity and specificity of state-dependent neural activity in the brainstem

To assess state-dependent firing of individual neurons in the brainstem on a timescale of seconds to minutes, we performed *in vivo* silicon probe recording (**Fig. 2A**) from 7 head-fixed mice (9 recording sessions) and examined how individual neurons change their firing across behavioral states. **Figure 2B** shows representative examples of state-dependent firing from four simultaneously recorded neurons. Even within a particular state from the same animal, brainstem neurons show highly diverse and dynamic firing.

To classify neurons according to their state-dependent firing, we computed mean firing rate in each state across neurons (n = 76) and applied a hierarchical clustering algorithm (**Fig. 2C**). We identified four functional classes: awake (AW)-on neurons (23.7 %) were more active during wakefulness compared to sleep states. REM-off neurons (17.1 %) reduced their firing during REM sleep. REM/AW-on neurons (6.6 %) were quiet during NREM sleep. The largest class (52.6 %) was REM-on neurons, which showed the highest firing rate during REM sleep. Thus, we confirmed highly diverse state-dependent firing in the brainstem.

Because the recorded neurons were distributed across various nuclei in the brainstem, it was difficult to determine their state-dependency in each nucleus. However, a subset of neurons was likely recorded from the cholinergic system, namely the pedunculopontine tegmental nucleus and the laterodorsal tegmental nucleus, which show AW-on or REM-on activity (**Supplementary Fig. 3**). To verify this, we performed *in vivo* fiber photometry of Ca^2+^ signals from pontine cholinergic neurons by expressing GCaMP6s in freely behaving mice (4 animals, 12 recording sessions) (**Fig. 2D**). Consistent with the data from *in vivo* electrophysiology, cholinergic populations showed larger activity during REM sleep and wakefulness, compared to NREM sleep (F_2,32_ = 5.12, *p* = 0.012, one-way ANOVA) (**Figs. 2E-G**). Therefore, although state-dependency of individual neuronal firing in the brainstem is diverse, we also confirmed state-dependent and cell-type-specific firing.

### Behavioral correlates of the sleep-wake cycle and underlying neural activity in the brainstem

Pupil diameter is an excellent biomarker of global brain state or arousal level (Aston-Jones and Cohen, 2005; Larsen and Waters, 2018; McGinley et al., 2015; Yuzgec et al., 2018) and activity in brainstem neurons, especially locus coeruleus norepinephrine neurons, correlates with pupil diameter (Aston-Jones and Cohen, 2005). However, it is still unclear how pupil dilation changes around the transition of sleep-wake states and to what extent brainstem neurons as a population can predict pupil dilation quantitatively. To address these issues, we analyzed datasets from head-fixed mice with either silicon probe recordings from the brainstem (6 animals, 6 recording sessions) or the hippocampus (2 animals, 3 recording sessions), or field potential recording from the brainstem with a bipolar electrode (6 animals, 9 recording sessions).

As previously reported (Yuzgec et al., 2018), mice in a head-fixed condition kept their eyes open, allowing us to monitor pupil dilation across states along with cortical EEG and EMG. The effects of behavioral states on pupil diameter was statistically significant (**Fig. 3A**, F_2, 53_ = 220.33, *p* < 0.0001, one-way ANOVA). More specifically, pupil diameter was constricted during REM sleep and dilated during wakefulness.

With respect to pupil dilation dynamics across states (**Fig. 3B and Supplementary Fig. 4**), pupil diameter dynamically fluctuated during wakefulness and gradually constricted during NREM sleep. Typically, 10-20 sec before REM sleep, the pupil diameter would further decrease and was fully constricted during REM sleep, with rapid eye movement (**Supplementary Fig. 4**).

Taking advantage of the simultaneous neural population recording and pupil monitoring, we examined how brainstem neurons can predict pupil dilation. First, we predicted pupil dilation based on the activity of individual neurons (**Fig. 3C**) by applying a linear regression analysis. Because it was expected that the preceding neural activity can better predict pupil dilation, we systematically shifted the temporal relationship between spike trains and pupil diameter (see Methods). As expected, most of the neurons showed asymmetric profiles of R^2^ values (**Fig. 3C**). Although individual profiles were diverse, the average profile showed predictive activity of brainstem neurons for pupil diameter around 10 seconds in advance. We also predicted pupil diameter based on simultaneously recorded brainstem neurons (**Fig. 3D**) by applying a multiple linear regression analysis. As with individual neurons, we observed an asymmetric profile of predictability. Thus, changes in brainstem neural activity preceded those in pupil diameter. Thereby, brainstem populations have predictive power for pupil diameter.

### Longer predictability of brainstem ensembles for vigilance states

Next, we examined whether and to what extent brainstem neurons have predictive power for behavioral states. To address this, first, we extracted features of neural population activity by applying non-negative matrix factorization (NMF) (Lee and Seung, 1999; Onken et al., 2016) (**Figs. 4A and B**). While overall firing rate reflected state changes (**Fig. 4A**), NMF could extract several modules which captured state-dependent firing patterns across neurons in an unsupervised fashion. For example, module 1 represented REM-on activity whereas module 2 was activated at the end of NREM sleep and module 3 was most active during wakefulness. Indeed, the weights in each module were consistent with state-dependency of individual neural firing (**Supplementary Fig 5**).

Besides modules capturing firing patterns across neurons, NMF also yielded activation coefficients of these modules. We noticed that dynamics of these activation coefficients show predictive activity: in the case of Figure 4A, the activation coefficients of modules 1 and 2 gradually built up during REM and NREM sleep, respectively. Therefore, we hypothesized that brainstem population activity exhibits not just state-dependency, but also predictive power for behavioral states (i.e., wakefulness, NREM sleep and REM sleep). To test this, we took the activation coefficients profiles from three modules and classified behavioral states by training a linear classifier, with systematic time shifting (**Fig. 4C**). Brainstem populations showed predictive activity 10s of seconds before transitions to all three behavioral states. To compare, we also performed the same analysis for hippocampal neural activity (**Fig. 4C**). Although hippocampal neurons also had predictive power for several tens of seconds, the profile was relatively short-lasting compared to that of brainstem neurons. Thus, brainstem neurons have long-lasting predictive power for behavioral states compared to hippocampal neurons.

### Brainstem population activity underlying P-waves during NREM and REM sleep

On a timescale of seconds to minutes, brainstem neurons show diverse but specific state-dependent firing and have predictive power for pupil dilation and behavioral states. To investigate brainstem neural firing on a sub-second timescale, we focused on P-waves (Callaway et al., 1987; Datta, 1997). Although these sub-second neural events in the brainstem have long been recognized, the underlying neural ensembles and relations to other sleep-related oscillations are fully understood.

Taking advantage of our dataset, we first examined whether the mouse pons exhibits P-waves like other mammalian species. We implanted a bipolar electrode in the pons (n = 16 recordings) (**Fig. 5A**) and monitored LFPs by subtracting signals. During REM sleep, we observed large amplitude irregular neural events, which often appeared as a burst (**Fig. 5B right**). We also observed similar, but isolated neural events during NREM sleep (**Fig. 5B left**). These neural events appeared more often during REM sleep (*p* < 0.0001, two-tailed *t*-test) (**Fig. 5C**). Intriguingly, the frequency of these events gradually increased during NREM to REM sleep transitions and decreased during REM sleep to wakefulness transitions (**Fig. 5D**). Because these characteristics generally resemble to those in other species (Callaway et al., 1987; Datta, 1997), we concluded that these neural events are P-waves in mice.

P-waves can also be seen in silicon probe recordings (**Fig. 5E**). Similar large-amplitude, irregular neural events were observed in subtracted and filtered LFPs (**Fig. 5F**). Many of the simultaneously recorded brainstem neurons fired during P-waves. To assess this tendency, we pooled the peri-event firing profiles of all recorded brainstem neurons around P-waves (**Figs. 5G and H**). The firing profiles were aligned at the trough timing of P-waves. A subset of neurons showed peak firing at the falling phase of P-waves. This tendency was consistent between NREM and REM sleep, suggesting that P-waves during NREM sleep (P-waves^NREM^) are equivalent to P-waves during REM sleep (P-waves^REM^), with respect to neural firing within the brainstem.

### State-dependent functional interactions between P-waves and cortical activity

Co-firing of a subset of brainstem neurons underlies P-waves during both NREM and REM sleep. What are the impacts of such neural events onto other brain regions? Are any other sleep-related neural events associated with P-waves? Addressing these questions would provide insight into functions of P-waves. To this end, first, we investigated the relationship between P-waves and cortical EEGs (**Fig. 6**). During NREM sleep, P-waves were associated with multiple oscillatory components. Averaged P-wave-triggered cortical EEGs exhibited multiple phasic components (**Fig. 6B**), which consisted of delta (1-4 Hz), theta (∼7 Hz) and beta (15∼30 Hz) frequencies. To examine this trend further, we assessed the phase relationship between P-wave timing and cortical oscillations (**Fig. 6C**). We found significant phase preferences of P-wave timing (*p* < 0.01, Rayleigh’s test). We further assessed this phase-locking activity by computing phase modulation index, which is defined as the difference in proportions of P-waves between the preferred phase and opposite phase by dividing phases into four bins (e.g., the higher phase modulation index reflects the larger difference in the proportion of P-waves between two opposing phase bins) (**Fig. 6D**). We found larger phase modulation at delta and beta ranges.

On the other hand, P-waves^REM^ exhibited distinct associations with cortical oscillations (**Figs. 6B-D**). We observed significant phase modulation at theta range (*p* < 0.01) (**Fig. 6C**), indicating that two prominent neural markers in REM sleep, that is, theta oscillations and P-waves, are temporally organized.

Next, we investigated the relationship between P-waves and hippocampal activity (**Fig. 7**). We started by assessing the phase relationship between hippocampal LFPs and P-waves across frequency bands (**Fig. 7B**). While the timing of P-waves was phase-locked strongly at theta range (∼7 Hz) in both sleep states, we also observed stronger phase modulations with high frequency components during REM sleep. We also examined underlying spiking activity in the hippocampus (**Fig. 7C**). Intriguingly, while a subset of hippocampal neurons fired most strongly around the timing of P-waves during both NREM and REM sleep, the temporal order between hippocampal neural firing and P-waves was state-dependent (**Fig. 7C**): during NREM sleep, co-firing of hippocampal neurons was followed by P-waves whereas P-waves were followed by burst firing in subset of hippocampal neurons during REM sleep. To test the hypothesis that co-firing of hippocampal neurons during NREM sleep may reflect sharp-wave ripples (SWRs), we detected high-frequency ripple events based on hippocampal LFPs to assess the temporal relationship between ripples and P-waves^NREM^ (**Fig. 7D**). We found that ripple events preceded P-waves during NREM sleep. Thus, P-waves are strongly associated with hippocampal activity in both sleep states. However, their associations are state-dependent.

## Discussion

Although state-dependent neural ensembles have been intensively characterized in the cortex, little is known about the brainstem. Here, we investigated state-dependent neural population activity in the brainstem, primarily the pons, on two distinct timescales. On a timescale of seconds to minutes, brainstem neurons show diverse state-dependent firing, with cell-type-specificity in pontine cholinergic neurons. Brainstem activity can collectively predict pupil dilation as well as behavioral states. The ability to predict behavioral states is longer lasting compared to hippocampal neurons. These relatively slow dynamics may be related to observations from optogenetic experiments where the effect of optogenetic stimulation on state transitions often emerges tens of seconds after stimulus onset (Adamantidis et al., 2007; Tsunematsu et al., 2013; Tsunematsu et al., 2014; Van Dort et al., 2015; Zhang et al., 2019).

On a timescale of sub-seconds, we characterized P-waves in the mouse, with respect to underlying neural firing as well as associated cortical activity. P-waves typically appear during REM sleep and less during NREM sleep. P-waves in both sleep states are accompanied by synchronous firing of brainstem neurons, suggesting that underlying local activity during P-waves is similar between sleep states. However, their relationship to cortical neural events is state-dependent: the timing of P-waves^NREM^ are phase-locked to various cortical oscillations and hippocampal SWRs precede P-waves^NREM^, suggesting that P-waves are part of the brain-wide neural events triggered by SWRs. During REM sleep, P-waves are phase-locked most strongly at theta frequency in both the neocortex and hippocampus. Crucially, P-waves precede firing in a subset of hippocampal neurons, suggesting that P-waves may trigger brain-wide neural events. Thus, P-waves are part of the state-dependent coordinated activity across the brain.

### Technical considerations

State-dependent activity in the brainstem has been described over the past several decades by using various methods. The present study utilized a silicon probe to monitor neural activity from multiple neurons simultaneously at a high temporal resolution. This approach allowed us to (1) quantify state dependency of brainstem neural ensemble dynamics on a timescale of seconds to minutes and (2) characterize neural population activity underlying P-waves for the first time. However, because silicon probe recording alone has a limitation to identify cell types, additional approaches, such as Ca^2+^ imaging (**Fig. 2**) or electrophysiology with optogenetic tagging (Weber et al., 2015; Yague et al., 2017; Zhang et al., 2019), can complement this study to determine how specific types of neurons contribute to state-dependent neural ensembles in the brainstem.

### Slow dynamics of brainstem ensemble dynamics

Our results in **Figures 3 and 4** are consistent with the notion that brainstem populations play a regulatory role in pupil dilation/constriction (Aston-Jones and Cohen, 2005; Larsen and Waters, 2018) as well as global brain states (Brown et al., 2012; Herice et al., 2019; Luppi et al., 2012; Weber and Dan, 2016). Crucially, the asymmetric profile of the predictability for pupil diameter suggests that the modulation of brainstem activity precedes pupil dilation, rather than simple correlations.

The long lasting predictability of brainstem populations for behavioral states is not trivial. Intriguingly, the slow (30-60 sec) timescale recalls us the timescale observed in some of optogenetic experiments: although optogenetic stimulation can modulate neural firing at a millisecond resolution, the effect of optical stimulation on state transitions typically emerges tens of seconds after stimulus onset (Adamantidis et al., 2007; Van Dort et al., 2015; Zhang et al., 2019). The exact mechanism is still unknown, but we hypothesize that the modulation of neural activity in the brainstem occurs tens of seconds before global brain state transitions from one state to another. In other words, each state emerges from complex interactions across various regions of the brain.

### P-waves in mice

Although PGO or P-waves have been studies since the 1970s in several mammalian species, to the best of our knowledge, we are the first to characterize P-waves in mice. Given the growing importance of the mouse as an animal model for sleep research (Herice et al., 2019), the confirmation of P-waves in mice is important for further interrogation.

We have noticed several similarities and differences in P-waves between mice and other species. First, the waveform of P-waves in mice is generally consistent with those in other species, such as cats (Callaway et al., 1987; Jeannerod et al., 1965) and rats (Datta, 1997; Farber et al., 1980), suggesting that underlying neural ensembles may be similar across species. Second, the frequency of P-waves during REM sleep is generally consistent across species (Datta, 1997). However, we have also noticed that the frequency of detected P-waves varied across our experiments. This may be explained by either the variation of REM sleep quality or the variation of electrode positions. Further analysis of P-waves across brainstem nuclei will provide insights into their relationship with sleep homeostasis and the mechanism of P-wave genesis. Third, the temporal evolution of P-wave frequency generally agrees between mice and cats: the frequency of PGO-waves gradually increases before the transition of NREM to REM sleep in cats (Steriade et al., 1989). Although it was weak, a similar tendency was observed in our recordings (**Fig. 5D**). Rather, the frequency of P-waves increases during REM sleep. This subtle difference may be explained by anatomical differences between species (Datta, 2012). Finally, although P-waves appear more frequently during REM sleep, it is important to note that similar neural events also occur during NREM sleep. Given state-dependent interactions between P-waves and cortical oscillations (**Figs. 6 and 7**), the mechanisms of P-wave genesis are likely distinct.

### State-dependent global coordination of neural ensembles

The temporal correlation between P-waves and hippocampal theta rhythms during REM sleep is consistent with previous studies in cats and rats (Karashima et al., 2004; Sakai et al., 1973). The phase-locked activity with fast gamma (80-110 Hz) in the hippocampus may relate to the recent observation that demonstrated the close association between local hippocampal theta and fast gamma events and brain-wide hyperemic events (Bergel et al., 2018). Because a number of hippocampal neurons fire immediately after P-waves (**Fig. 7C**), P-waves may play a role in the regulation of hippocampal ensemble dynamics as well as these brain-wide events during REM sleep.

On the other hand, the picture during NREM sleep seems to be distinct. Because SWRs precedes P-waves (**Fig. 7D**) and SWRs are known to be generated within hippocampal circuits (Buzsaki, 2015), SWRs play a leading role in brain-wide sub-second neural events including P-waves. These state-dependent brain-wide neural ensembles imply distinct functional roles of sleep states.

## Methods

### Animals

All experimental procedures were performed in accordance with the United Kingdom Animals (Scientific Procedures) Act of 1986 Home Office regulations and approved by the Home Office (PPL 70/8883). A total of 21 mice were used in this study (**Supplementary Table 1**). Their genotypes consisted of wild-type, ChAT-IRES-Cre (JAX006410), or ChAT-IRES-Cre::Ai32 (JAX012569) on C57BL/6 background. ChAT-IRES-Cre::Ai32 mice were used to identify pontine cholinergic neurons post-hoc histological analysis. For brainstem silicon probe recordings, 6 animals were used (9 recordings). For hippocampal silicon probe recordings, 2 animals were used (4 recordings). For P-wave recordings, 10 animals were used, but 4 were excluded due to electrode mispositioning or lack of histological data. 16 datasets were used for further analysis. For pupil monitoring, 14 animals were used, but one animal was excluded due to their eye closure during recording. 18 datasets were used for further analysis. For fiber photometry, 4 animals were used (12 recordings). The detailed information of genotypes, age, sex, body weight and the number of recordings was provided in **Supplementary Table 1**.

### Surgical procedures

For all *in vivo* electrophysiological experiments, mice were anesthetized with isoflurane (5% for induction, 1-2% for maintenance) and placed in a stereotaxic apparatus (SR-5M-HT, Narishige). Body temperature was maintained at 37°C with a feedback temperature controller (40–90–8C, FHC). Lidocaine (2%, 0.1-0.3 mg) was administered subcutaneously at the site of incision. Two bone screws were implanted on the skull as electrodes for cortical EEGs and twisted wires were inserted into the neck muscle as electrodes for EMG. Another bone screw was implanted on the cerebellum as a ground/reference.

For pontine EEG recording, bipolar electrodes were bilaterally implanted in the medial parabrachial nucleus of the pons (5.1 mm posterior, 1.2 mm lateral from bregma, 3.2 mm depth from brain surface). The bipolar electrodes consisted of 75 or 100 µm diameter stainless wires (FE631309, Advent Research Materials and FE205850, Goodfellow, respectively). The tip of two glued wires were separated by 0.5-1.0 mm vertically to differentiate EEG signals. All electrodes were connected to connectors (SS-132-T-2-N, Semtec) and securely attached on the skull with dental cement. A pair of nuts was also attached on the skull with dental cement as a head-post. After the surgery, Carprofen (Rimadyl, 5 mg/kg) was administered intraperitoneously.

For brainstem or hippocampal silicon probe recording, in addition to bone screws for cortical EEGs and a ground/reference, a pair of nuts was attached on the skull with dental cement as a head-post. After the head-post surgery, the animals were left to recover for at least 5 days. During the habituation period, the animals were placed in a head-fixed apparatus, by securing them by the head-post and placing the animal into an acrylic tube. This procedure was continued for at least 5 days, during which the duration of head-fixed was gradually extended from 10 to 120 min. A day after the habituation period, the animals were anesthetized with isoflurane and a craniotomy to insert silicon probe the brainstem and hippocampus was performed. A craniotomy on the left hemisphere (4.0 mm to 5.5 mm posterior, 1.0 to 1.3 mm lateral from bregma) for the brainstem recording and on the left hemisphere (2.0 mm posterior, 1.5 mm lateral from bregma) for the hippocampus recording were performed, respectively. To protect and prevent the brain from drying, the surface was covered with biocompatible sealants (Kwik-Sil and Kwik-Cast, WPI). In the following day, the animals were placed in the head-fixed apparatus for electrophysiological recording as described below.

For fiber photometry experiments, cortical EEG and EMG electrodes were implanted as described above and connected to a 2-by-3 piece connector (SLD-112-T-12, Semtec). Two additional anchor screws were implanted bilaterally over the parietal bone to provide stability and a small portion of a drinking straw was placed horizontally between the anchor screws and the connector. The viral vector (AAV5-CAG-flex-GCaMP6s-WPRE-SV40, Penn Vector Core; titer 8.3×10^12^ GC/ml) was microinjected (500 nl at 30 ml/min) (Nanoliter2010, WPI) to target the PPT/LDT area (−4.5 mm posterior, 1 mm lateral from bregma and 3.25 mm depth from brain surface). The micropipette was left in the brain for an additional 10 minutes and then slowly raised up. An optic fiber cannula (CFM14L05, Thorlabs) was then implanted 3 mm deep from the surface of the brain and all components were secured to each other and the skull with dental cement.

### *in vivo* electrophysiological experiments in a head-fixed condition

Experimental procedures were as described previously (Lyngholm and Sakata, 2019; Yague et al., 2017). Briefly, all electrophysiological recordings were performed in a single-walled acoustic chamber (MAC-3, IAC Acoustics) with the interior covered with 3 inches of acoustic absorption foam. For pontine EEG recording, cortical EEG, EMG and pontine EEG signals were amplified (HST/32V-G20 and PBX3, Plexon), filtered (0.1 Hz low cut), digitized at a sampling rate of 1 kHz and recorded using LabVIEW software (National Instruments). Recording was performed for 5 hrs from 9:00 to 14:00. For brainstem or hippocampal silicon probe recording, a 32 channels 4 shank silicon probe (A4 × 8-5 mm-100-400-177 for brainstem recording or Buzsaki32 for hippocampus recording) was inserted slowly with a manual micromanipulator (SM-25A, Narishige) into the brainstem (3.75 mm – 4.3 mm from the brain surface) or the hippocampus (1.55 mm – 1.85 mm from brain surface). Probes were inserted perpendicularly with respect to the brain surface. Broadband signals were amplified (HST/32V-G20 and PBX3, Plexon) relative to screw on the cerebellum, filtered (0.1 Hz low cut), digitized at 20 kHz and recorded using LabVIEW software (National Instruments). The recording session was initiated > 1 hr after the probe was inserted to its target depth, to stabilize neural signals. Recording preparation started from 8:00 and terminated by 15:00. For verification of silicon probe tracks, the rear of the probes were painted with DiI (∼10% in ethanol, D282, Invitrogen) before insertion.

### Pupil monitoring

In a subset of *in vivo* electrophysiological experiments under a head-fixed condition, pupil was also monitored with an off-axis infrared (IR) light source (860 nm IR LED, RS Components). A camera (acA1920-25µm, Basler Ace) with a zoom lens (M0814-MP2, computar) and an IR filter (FGL780, Thorlabs) was placed at ∼10 cm from the animal’s left eye. Images were collected at 25 Hz using a custom-written LabVIEW program and a National Instruments image grabber (PCIe-8242).

### *in vivo* fiber photometry experiments in a freely behaving condition

The fiber photometry system consisted of two excitation channels. A 470 nm LED (M470L3, Thorlabs) was used to extract a Ca^2+^-dependent signal and a 405 nm LED (M405L3, Thorlabs) was used to obtain a Ca^2+^-independent isosbestic signal. Light from the LEDs was directed through excitation filters (FB470-10, FB405-10, Thorlabs) and a dichroic mirror to the fiber launch (DMLP425R and KT110/M, respectively), The fiber launch was connected to a multimode patch cable (M82L01, Thorlabs) which attached to an implantable optic fiber on the mouse via a ceramic mating sleeve (CFM14L05 and ADAF1, respectively). Light emissions from GCaMP6s expressing neurons were then collected back through the optic fiber, and directed through a detection path, passing a dichroic mirror (MD498) to reach a photodetector (NewFocus 2151, Newport). A National Instruments DAQ (NI USB-6211) and custom-written LabVIEW software was used to control the LEDs and acquire fluorescence data at 1 KHz. LEDs were alternately turned on and off at 40Hz in a square pulse pattern. Electrophysiology signals were recorded at 1 KHz using an interface board (RHD2000, Intan Technologies) and connected to the mouse via an amplifier (RHD2132, Intan Technologies). Mice were habituated to being handled and tethered to the freely behaving system over several consecutive days. Mice were scruffed and the straw on the headcap slotted into a custom-made clamp, to keep the head still and absorb any vertical forces when connecting the electrophysiology and fibre photometry tethers to the headcap. Once connected, mice were placed in an open top Perspex box (21.5 cm x 47 cm x 20 cm depth) lined with absorbent paper, bedding and some baby food. Recordings lasted 4-5 hours to allow for multiple sleep/wake transitions.

### Histological analysis

After electrophysiological experiments, animals were deeply anesthetized with mixture of pentobarbital and lidocaine and perfused transcardially with 20 mL saline followed by 20 mL 4% paraformaldehyde/0.1 M phosphatase buffer, pH 7.4. The brains were removed and immersed in the above fixative solution overnight at 4°C and then immersed in a 30% sucrose in phosphate buffer saline (PBS) for at least 2 days. The brains were quickly frozen and were cut into coronal or sagittal sections with a sliding microtome (SM2010R, Leica) or with a cryostat (CM3050, Leica) at a thickness of 50 or 100 µm. The brain sections were incubated with a NeuroTrace 500/525 Green-Fluorescence (1/350, Invitrogen) or NeuroTrace 435/455 Blue-Fluorescence (1/100, Invitrogen) as Nissl staining in PBS for 20 min at room temperature (RT) followed by incubating with a blocking solution (10% normal goat serum, NGS, in 0.3% Triton X in PBS, PBST) for 1 hr at RT. For ChAT-IRES-Cre::Ai32 mice, to confirm the position of ChAT-expressing neurons, we performed GFP and ChAT double staining. These brain sections were incubated with mouse anti-GFP antiserum (1/2000, ABCAM) and goat anti-ChAT antiserum (1/1000, Millipore) in 3% NGS in PBST for overnight at 4°C. After washing, sections were incubated with DyLight 488-labeled donkey anti-mouse IgG (1/500, Invitrogen) and Alexa 568-labeled donkey anti-goat IgG (1/500, Invitrogen) for 2 hrs at RT. After staining, sections were mounted on gelatin-coated or MAS-coated slides and cover-slipped with 50% glycerol in PBS. The sections were examined with a fluorescence microscope (BZ-9000, Keyence).

### Data analysis

#### Sleep scoring

Vigilance states were visually scored offline according to standard criteria (Radulovacki et al., 1984; Tobler et al., 1997). Wakefulness, NREM sleep, or REM sleep was determined in every 4 second based on cortical EEG and EMG signals using custom-made MATLAB GUI. For electrophysiological or fiber photometry experiments, the same individual scored all recordings for consistency.

#### Pupil analysis

Video files were processed by using DeepLabCut (Mathis et al., 2018). Each video file consisted of a 5 min segment of the recording, meaning that each experiment yields tens of video files. For training, a single file was chosen so that all states (AW, NREM sleep and REM sleep) could appear based on the sleep score. This initial step was critical to reflect the dynamic range of pupil dilation/constriction in each experiment. After choosing the file, 20 frames were randomly selected to manually mark the left and right edges of pupil. ImageJ was used for this manual marking. Using these labeled frames, the deep convolutional neural network was then trained, and all video files were processed to detect the left and right edges of pupil. After processing, visual inspection was performed by generating a down-sampled (50-100 times) video clip. The same procedure was applied across all recordings. To compute pupil diameter, the distance between the left and right edges of the pupil was calculated across frames. The profile was filtered (low-pass filter at 0.5 Hz) and z-scored. To compute eye movement, the middle point of pupil was determined and the distance of the middle points between two continuous frames was calculated. The profile was then normalized by the maximal value of the profile. From 20 pupil recordings (14 animals), 2 recordings from a single animal were excluded due to eye closure during most of recording period (**Supplementary Table 1**).

#### Fiber photometry signal processing

Custom-written MATLAB scripts were used to compute dF/F signals. To extract 405 and 470 nm signals, illumination periods were determined by detecting synchronization pulses. For each illumination epoch, the median fluorescent signal was calculated. Because each illumination epoch consisted of pulses at 40 Hz, the fluorescent signals originally sampled at 1 kHz were effectively down-sampled to 40 Hz. Photobleaching was estimated by a single exponential curve and the difference between the fluorescent signal trace and the estimate was further low-pass filtered at 4 Hz. To estimate moving artifacts, the filtered 405 nm signals were linearly scaled based on the filtered 470 nm signals using a linear regression. To estimate dF/F signals, then the 470 nm signals were subtracted from the scaled 405 nm signals. In this study, the first 10 min segment was excluded for further analysis due to poor estimation of the photobleaching profile.

#### Spike train and LFP/EEG analysis

For spike sorting, Kilosort (Pachitariu et al., 2016) was used for automatic processing, followed by manual curation using phy (https://github.com/cortex-lab/phy). Clusters with ≥ 20 isolation distance were recognized as single units. The position of single units was estimated based on the channel providing the largest spike amplitude. All subsequent analysis were performed by using custom-written codes (MATLAB, Mathworks).

To categorize functional classes of single units, average firing rate for each behavioral state was calculated and a hierarchical clustering approach with the Ward’s method was applied.

To predict the pupil diameter from single unit activity, spike trains were filtered (band-pass filter between 0.5 and 25 Hz) and then a liner regression analysis was performed. To evaluate the goodness-of-fit of the linear model, R-squared value was calculated. The same process was repeated by shifting the time relationship between spike trains and pupil diameter to determine an optimal time window to predict pupil diameter from spike train. Then the sequence of R-squared values were normalized by computing Z-scores.

To predict the pupil diameter from simultaneously monitored multiple single unit activities, spike trains were filtered (band-pass filter between 0.5 and 25 Hz) and a linear regression model was trained by using a regularized support-vector machine algorithm with 10-hold cross-validation. Then cross-validated mean squared error (MSE) was computed. The same process was repeated by shifting the time relationship between spike trains and pupil diameter. The sequence of MSEs were normalized.

To decompose neural population activity into “space” (neurons) modules and activation coefficients of these modules, we adopted non-negative matrix factorization (NMF) (Lee and Seung, 1999; Onken et al., 2016). First, spike trains were discretized by binning them into 4 sec intervals, which were equivalent to the time window for sleep scoring (see above). Let **r**(*t*) denote the resulting vector of population spike counts in bin *t*. We represented all population spike count vectors in a matrix **R** = [**r**(1) **r**(2) … **r**(*T*)] of size *N* by *T*, where *N* denotes the number of neurons and *T* denotes the number of time bins. We then decomposed the matrix **R** into two non-negative matrices **W** of size *N* by *m* and **H** of size *m* by *T* as follows: **R** = **WH**, where *m* is the number of modules. To this end, we applied multiplicative update rules (Lee and Seung, 2001):

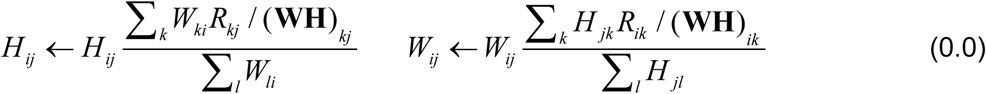

These update rules minimize the Kullback-Leibler divergence, corresponding to a Poisson noise assumption for the spike counts (Févotte and Cemgil, 2009). In each run, we randomly initialized **W** and **H** and applied the update rules up to 100 times or until convergence. For each decomposition, we performed 10 runs and selected the run with the lowest Kullback-Leibler divergence. The *m* columns of **W** then represented the *m* space modules and the *m* corresponding rows of **H** represented their activation coefficients for each time bin.

To select the number of space modules *m*, we evaluated how many modules were needed so that additional modules did not significantly improve the decomposition. Our procedure was similar to that used in De Marchis et al. (De Marchis et al., 2013). We generated surrogate data by shuffling all elements of the matrix **R** and then decomposed the shuffled matrix like we previously decomposed the original **R**. We quantified the quality of a decomposition using the variance accounted for (VAF) (Clark et al., 2010). Starting with one module, we incremented the number of modules until the VAF of the unshuffled data decomposition did not exceed 3/4 of the average VAF of the decompositions of 100 shuffles.

To classify three behavioral states based on the activation coefficients, a linear classifier was trained by fitting a multivariate normal density to each state with 10-fold cross validation. Then classification performance was calculated. This procedure was repeated by shifting the time relationship between the activation coefficients and sleep scores.

To detect P-waves, two EEG or LFP signals in the pons were subtracted and filtered (5-30 Hz band-pass filter). If the signals cross a threshold, the event was recognized as P-waves. To determine the detection threshold, a 1-min segment of the subtracted signals was extracted from the longest NREM sleep episode for a reliable estimation of stable noise level. The noise level was estimated by computing root-mean-square (RMS) values in every 10 ms time window. The threshold was defined as mean + 5 x standard deviation of the RMS values. The timing of P-waves was defined as the timing of the negative peak.

The phase analysis was essentially the same as that described previously (Yague et al., 2017). Cortical EEG or hippocampal LFP signals were used for this analysis. For hippocampal LFPs, signals from two separate channels were subtracted to minimize volume conduction. To derive band-limited signals in different frequency bands, a Kaiser finite impulse response filter was used with sharp transition bandwidth of 1 Hz, pass-band ripple of 0.01 dB and stop-band attenuation of 50 dB. For filtering, MATLAB ‘filtfilt’ function was used. In the present study, the following bands were assessed: [2-4], [4-7], [7-10], [10-15], [15-20], [20-30], [30-50], [50-80], [80-110], and [110-150] Hz. The instantaneous phase of each band was estimated from the Hilbert transform and the phase of P-wave occurrence was computed. To quantify the relationship between P-waves and EEG/LFP phase, the percentage of P-waves elicited in each phase bin was calculated. The phase modulation was defined as the percentage in the preferred bin (the bin with maximal percentage) minus that in the opposite bin (the bin 180° apart). Rayleigh’s test for non-uniformity of circular data was performed to assess the statistical significance (*p* < 0.01) of the non-uniformity of the P-wave vs EEG/LFP phase distribution using CircStats Toolbox (Berens, 2009).

To detect ripples in the hippocampus, LFP signals from the channel which detected spiking activity were used. Band-limited signals at 140-250 Hz were computed by using a Kaiser finite impulse response filter (see above). Two sequences of RMS values were calculated with two different time window: 2 sec (long RMS) and 8 ms (short RMS). If the short RMS exceeds 4 times larger long RMS for 8 ms, then signals were recognized as a ripple event.

### Statistical analysis

Data was presented as mean ± SEM unless otherwise stated. Statistical analyses were performed with MATLAB. In **Figs. 2G and 3A**, one-way ANOVA with *post-hoc* Tukey’s Honest Significant Difference (HSD) criterion was performed. In **Fig. 4C**, repeated measures ANOVA was performed. In **Fig. 5C**, two-tailed *t*-test was performed. In **Fig. 6C**, Rayleigh’s test for non-uniformity was performed. To estimate effect size, Hedges’ *g* was computed using Measures of Effect Size Toolbox (Hentschke and Stuttgen, 2011).

## Acknowledgements

This work was supported by BBSRC (BB/M00905X/1), Leverhulme Trust (RPG-2015-377), Alzheimer’s Research UK (ARUK-PPG2017B-005), and Action on Hearing Loss (S45) to SS, EPSRC (EP/S005692/1) to AO, and a JSPS Postdoctoral Fellowship for Research Abroad, Research Fellowship from the Uehara Memorial Foundation, PRESTO from JST and KAKENHI (17H06520) to TT.

## Author contributions

TT performed all *in vivo* electrophysiological experiments and associated histological analysis and sleep scoring. AAP performed fiber photometry experiments and associated histological analysis and sleep scoring. SS performed all other data analysis. AO contributed to NMF analysis. TT, AAP, AO, and SS wrote the manuscript.

## Declaration of Interests

The authors declare no competing interests.

