## Supplemental Information for "State-dependent pontine ensemble dynamics and interactions with cortex across sleep states"

A

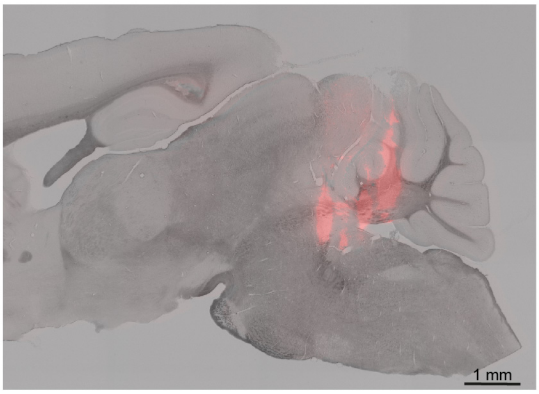


B

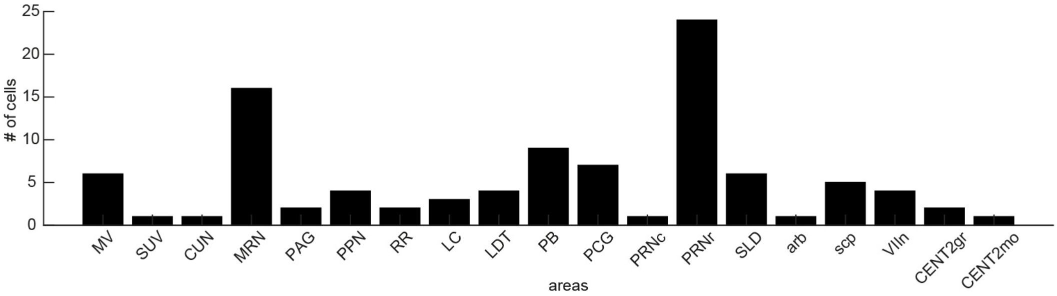


**Supplementary Figure 1. Histological analysis.**

(A) A sagittal section with showing the track of a silicon probe.

(B) The number of neurons recorded from brainstem nuclei in this study. MV, medial vestibular nucleus; SUV, superior vestibular nucleus; CUN, cuneiform nucleus ; MRN, midbrain reticular nucleus; PAG, periaqueductal gray; PPN, pedunculopontine nucleus; RR, retrorubral area; LC, locus ceruleus; LDT, laterodorsal tegmental nucleus; PB, parabrachial nucleus; PCG, pontine central gray; PRNc, pontine reticular nucleus, caudal part; PRNr, pontine reticular nucleus, rostral part; SLD, sublaterodorsal nucleus; arb, arbor vitae; scp, superior cerebellar peduncles; VIIn, facial nerve; CENT2gr, lobule II, granular layer; CENT2mo,lobule II, molecular layer.

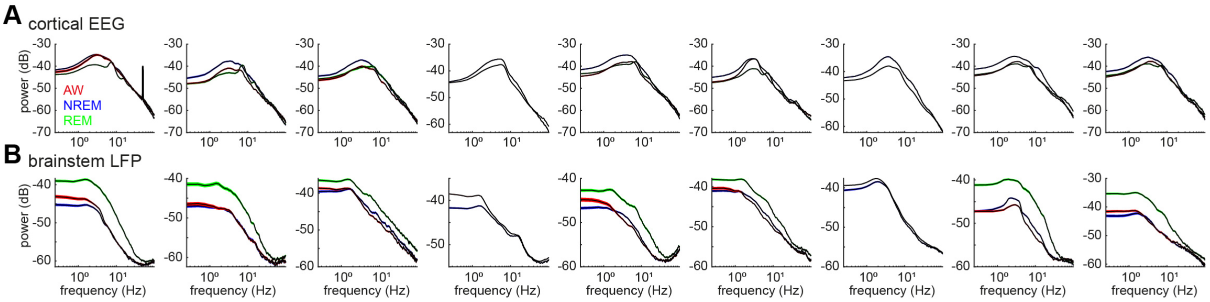


**Supplementary Figure 2. Power spectrum density of cortical EEGs (A) and brainstem LFPs (B) across all recording sessions.** Spectrum was computed in every 4-sec window in each state.

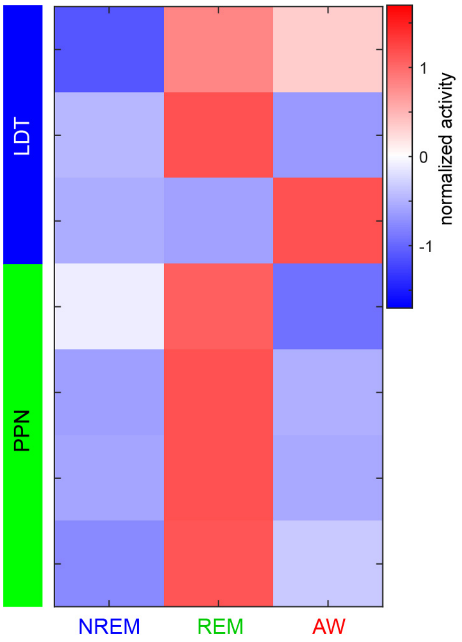


**Supplementary Figure 3. Neurons recorded from pontine cholinergic nuclei and their state dependency.**

Firing rates across three behavioral states were normalized as z-score. All neurons were classified as either REM-on, AW-on or REM/AW-on neurons.

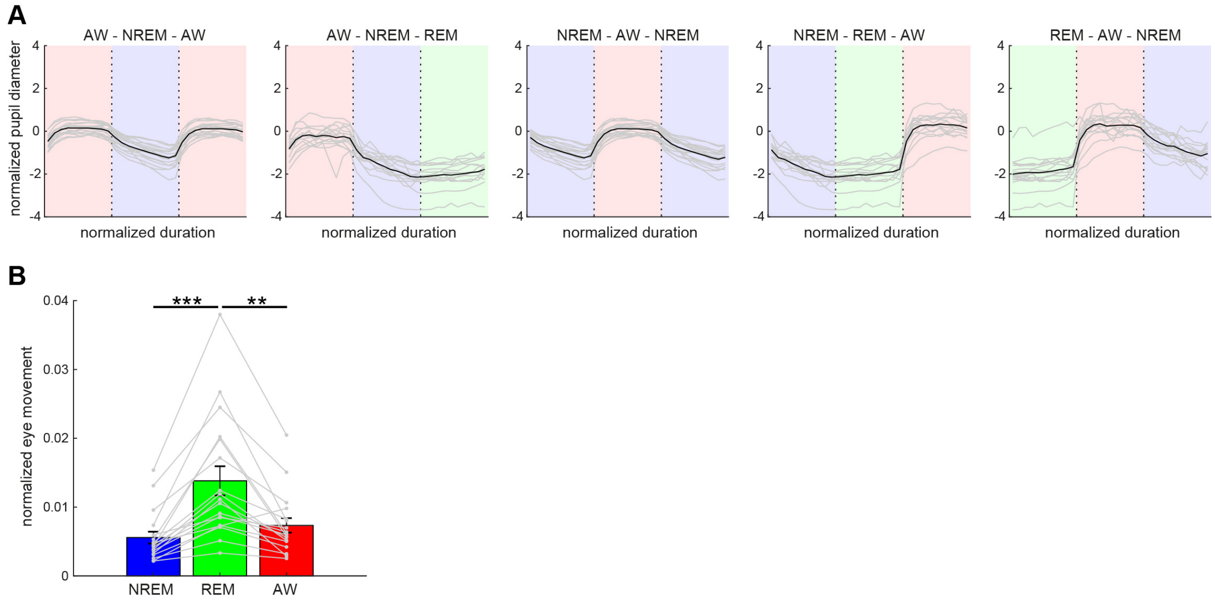


**Supplementary Figure 4. Pupil dilation and eye movement across vigilance states.**

(**A**) Normalized (z-scored) pupil diameter across state transitions. The duration of each state episode was normalized. Pupil diameter gradually decreased before REM sleep and stayed fully constricted during REM sleep.

(**B**) The average of eye movement across states. The effect of states on eye movement was significant ($F_{2,53}$ = 8.93, $p$ < 0.001, one-way ANOVA with post-hoc Tukey's honest significant difference criterion). **, p < 0.01; *** p < 0.001.

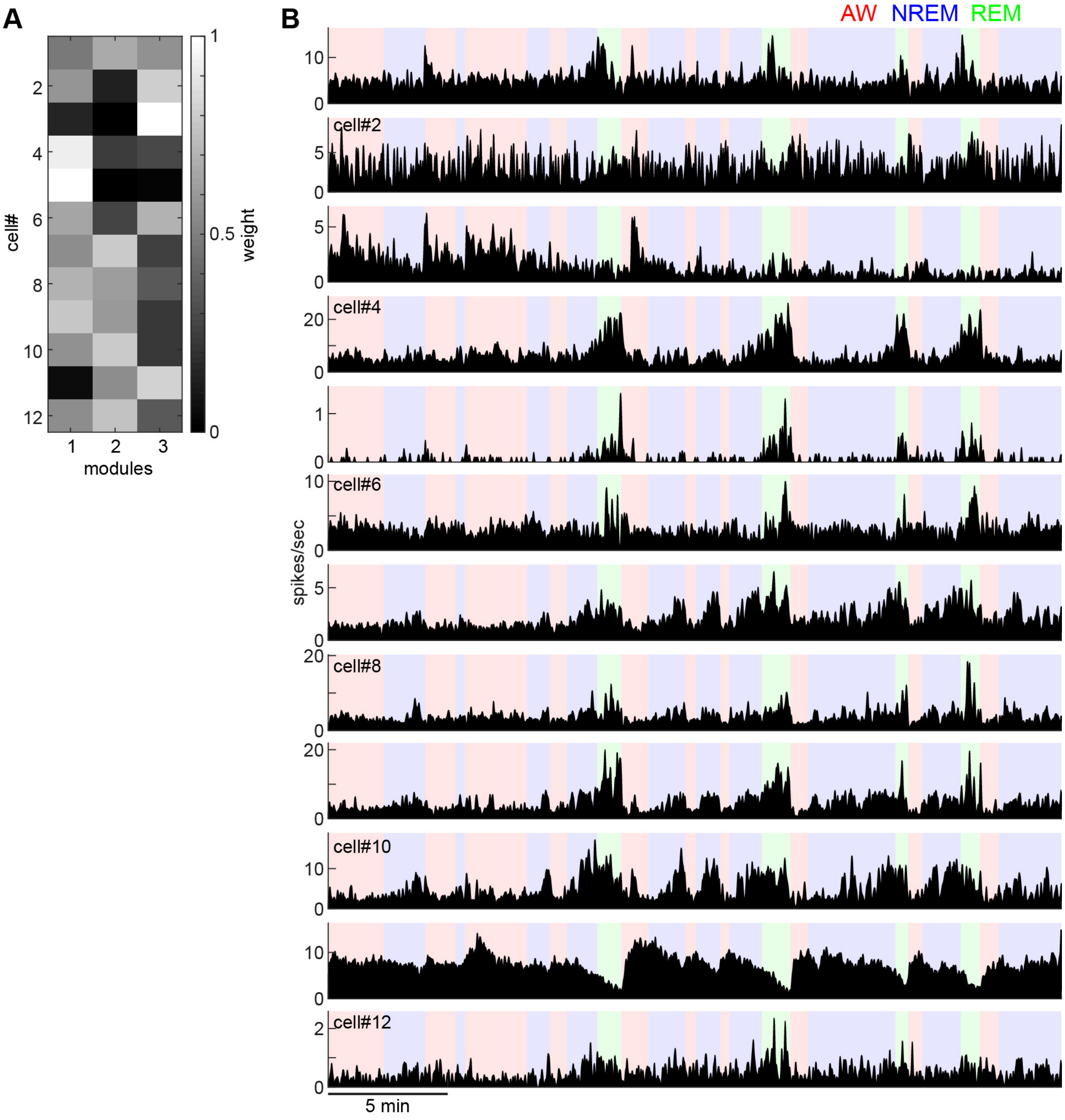


**Supplementary Figure 5. Weights of each module derived from non-negative matrix factorization (A) and individual firing profiles (B).**

Neurons with similar state-dependent firing patterns tend to have similar weights in each module. For example, cell#4 and #5 fire more during REM sleep and had higher weights in the module 1.

| Animal ID | Genotype | Sex | Weight (g) | Age (wk) | Experiment Type | # of recordings | | | | |
|---|---|---|---|---|---|---|---|---|---|---|
| | | | | | | BS-Si | BS-EEG | HP-Si | FP | Pupil |
| 1 | ChAT-Cre::Ai32 | M | 26.9 | 13 | BS-Si | 1 | | | | 1 |
| 2 | ChAT-Cre::Ai32 | M | 29.1 | 13 | BS-Si | 1 | | | | 1 |
| 3 | ChAT-Cre::Ai32 | M | 27.3 | 13 | BS-Si | 2 | | | | Exc[3] |
| 4 | ChAT-Cre::Ai32 | M | 34.6 | 26 | BS-Si | 1 | | | | 1 |
| 5 | WT | M | 28.0 | 14 | BS-Si | 1 | | | | 1 |
| 6 | WT | M | 24.7 | 11 | BS-Si | 2 | | | | 1 |
| 7 | WT | M | 27.0 | 11 | BS-Si | 1 | | | | 1 |
| 8 | ChAT-Cre::Ai32 | M | 30.7 | 19 | HP-Si & BS-EEG | | 1 | 2 | | 3 |
| 9 | ChAT-Cre::Ai32 | M | 33.4 | 19 | HP-Si & BS-EEG | | 2 | 2 | | 1 |
| 10 | ChAT-Cre | M | 28.6 | 16 | BS-EEG | | 3 | | | 3 |
| 11 | WT | M | 23.2 | 8 | BS-EEG | | 2 | | | NA |
| 12 | WT | M | 25.8 | 8 | BS-EEG | | 4 | | | NA |
| 13 | WT | M | 26.1 | 8 | BS-EEG | | 4 | | | NA |
| 14 | ChAT-Cre::Ai32 | M | 27.2 | 13 | BS-EEG | | Exc[1] | | | 2 |
| 15 | ChAT-Cre::Ai32 | M | 30.2 | 13 | BS-EEG | | Exc[2] | | | 1 |
| 16 | ChAT-Cre::Ai32 | M | 31.5 | 26 | BS-EEG | | Exc[2] | | | 1 |
| 17 | ChAT-Cre | M | 31.2 | 19 | BS-EEG | | Exc[2] | | | 1 |
| 18 | ChAT-Cre | F | 25.3 | 17 | Fiber photometry | | | | 3 | NA |
| 19 | ChAT-Cre | M | 25.5 | 12 | Fiber photometry | | | | 4 | NA |
| 20 | ChAT-Cre | M | 30.1 | 18 | Fiber photometry | | | | 3 | NA |
| 21 | ChAT-Cre | F | 20.4 | 21 | Fiber photometry | | | | 2 | NA |

**Supplementary Table 1. Summary of experimental animals and recordings in this study.**

BS-Si, silicon probe recording in the brainstem. HP-Si, silicon probe recording in the hippocampus. BS-EEG, EEG recording in the brainstem. FP, fiber photometry. Exc[1], the animal was excluded due to electrode mispositioning. Exc[2], the animal was excluded due to lack of histological data. Exc[3], the animal was excluded due to eye closure during recording.
